# Target capture sequencing provides insights into hybridogenetic water frogs

**DOI:** 10.1101/2025.09.18.677054

**Authors:** Lisanne van Veldhuijzen, Maciej Pabijan, Tariq Stark, Richard P.J.H. Struijk, Ben Wielstra, James France

**Affiliations:** Institute of Biology Leiden, Leiden University, P.O. Box 9505, 2300 RA Leiden, The Netherlands; Naturalis Biodiversity Center, P.O. Box 9517, 2300 RA Leiden, The Netherlands; Department of Comparative Anatomy, Institute of Zoology and Biomedical Research, Faculty of Biology, Jagiellonian University, ul. Gronostajowa 9, 30-387 Kraków, Poland; Reptile, Amphibian and Fish Conservation Netherlands (RAVON), P.O. Box 1413, 6501 BK Nijmegen, The Netherlands

**Keywords:** Amphibians, Sequence capture, Triploid, Hybridization

## Abstract

The European water frogs of the genus *Pelophylax*, with their propensity to hybridize and a non-orthodox inheritance system in some hybrids, present a genetic puzzle that has previously lacked strong genomic tools. This has limited investigation into the remarkable hybridogenetic system observed in the edible frog *P. esculentus* as well as inhibited monitoring of invasive populations in a genus that is critically vulnerable to this threat. In this study we explore whether a previously developed target capture bait set, FrogCap, can be used to answer genomic questions within the genus, by applying it to the three taxa of *Pelophylax* native to the Netherlands. We observe that the target capture data cleanly separates the three taxa, confirming significant introgression of *P. lessonae* mtDNA into *P. ridibundus* from the Netherlands. We also show that target capture is an efficient method for identifying polyploidy in *P. esculentus* and is effective at determining ancestral contribution within our sample population. Targeted sequence capture using FrogCap is a useful tool to unravel the intricate evolution of *Pelophylax* water frogs.

## Introduction

The edible frog, *Pelophylax esculentus*, is one of the most prominent examples of an extraordinary mode of reproduction known as hybridogenesis (Dufresnes & Mazepa, 2020; Lavanchy & Schwander, 2019). *Pelophylax esculentus* is the fertile hybrid of the pool frog, *P. lessonae*, and the marsh frog, *P. ridibundus*, which are sympatric across a large region of northern Europe (Berger, 1967; Dufresnes & Mazepa, 2020). While *P. esculentus* carries the genomes of both of its parents, these do not undergo recombination with each other and the hybrid will only transmit the genome of one species to its offspring (unless the hybrid is polyploid) (Graf & Polls Pelaz, 1989; Uzzell et al., 1976; Uzzell & Berger, 1975). Generally, the same parental genome is transmitted for many generations, resulting in a clonally transmitted lineage that reproduces itself at the expense of a sexual host species. This behaviour results in *P. esculentus* being designated as a klepton (and sometimes abbreviated as *P*. kl. *esculentus*) (Dubois & Günther, 1982).

Due to the lack of recombination between the parental genomes, all *P. esculentus* are effectively F_1_ hybrids, even though they may be descended from many generations of backcrossing. Consequently, *P. esculentus* individuals share a similar set of characteristics, intermediate between the parental species, and so may be considered as a distinct taxon. The morphology of *P. lessonae* and *P. ridibundus* varies considerably across their range, making distinguishing the three taxa difficult in some areas (Arntzen, 1981; Meilink et al., 2024), however they retain distinct ecological preferences and mating calls (Roesli & Reyer, 2000).

The genome that is transmitted varies between populations. In the most well-known example, *P. esculentus* occurs in mixed populations with *P. lessonae*, referred to as the *lessonae-esculentus* (or L-E system), where *P. esculentus* selectively transmits only the *P. ridibundus* genome. In other populations the situation is essentially reversed to form the *ridibundus-esculentus* (R-E) system (Hoffmann et al., 2015; Holsbeek & Jooris, 2010; Uzzell & Berger, 1975). In either case, *P. esculentus* could theoretically mate with itself to regenerate the alternative parental species. However, the offspring of matings between two *P. esculentus* are rarely viable (Berger, 1967; Guex et al., 2002; Vorburger, 2001). This is presumed to be due to such offspring inheriting two copies of the clonal genome. As this genome never undergoes recombination, it is unable to efficiently purge deleterious recessive alleles, leading to the phenomenon known as Muller’s ratchet (Muller, 1964). In cases where two *P. esculentus* from different clonal lineages mate, offspring viability is notably higher (Guex et al., 2002; Vorburger, 2001).

In addition to the L-E and R-E systems, populations exist that include all three taxa (the L-E-R system), as well as populations containing solely *P. esculentus* (the E-E system) (Hoffmann et al., 2015). The existence of the EE system is associated with polyploid individuals, which also allow for recombination within the clonal genome. Furthermore, these systems also involve the production of unreduced gametes, which transmit both parental genomes, preserving hybridity (Christiansen, 2009; Christiansen & Reyer, 2009; Dubey et al., 2019). Polyploidy is not exclusive to the EE system and may be observed across the range of *P. esculentus* (Biriuk et al., 2016; Blommers-Schlösser, 1990; Christiansen & Reyer, 2009; Dedukh et al., 2022).

A significant body of research has focused on the cytology of hybridogenesis in *P. esculentus*. It is observed that within the gonocytes of tadpoles, the chromosomes forming one of the parental genomes are corralled into micronuclei and gradually degraded, whereas the other parental genome is duplicated prior to meiosis (Chmielewska et al., 2018; Dedukh et al., 2020). However, there has been comparatively little study of the genomics of the hybridogenetic system. Molecular investigation in *Pelophylax* has largely focussed on allozymes (Hotz et al., 2013; Mezhzherin et al., 2024), and studies of limited numbers of nuclear and mitochondrial markers (Dufresnes, Monod-Broca, et al., 2024; Papežík et al., 2024; Patrelle et al., 2011; Sagonas et al., 2020). A small number of RAD-sequencing based studies provide wider genomic data (Doniol-Valcroze et al., 2021; Dubey et al., 2019; Dufresnes & Dubey, 2020), but as RAD-sequencing cannot be targeted to specific loci (e.g. coding genes), the resulting datasets are difficult to integrate due to frequent gaps and non-overlapping marker sets.

The lack of genomic data presents a major barrier to understanding the evolution of hybridogenesis in *Pelophylax* and the molecular mechanisms behind it. Presumably genetic or epigenetic factors determine whether the *lessonae* or *ridibundus* genome is transmitted, however no candidate loci are known. The hybridogenetic system should exclude the possibility of nuclear DNA transfer between the parental species (as one of the genomes is eliminated prior to miosis). However, the system has been observed to be subject to a degree of lability (Biriuk et al., 2016; Dedukh et al., 2019) and allozyme and AFLP data suggest that nuclear introgression between parental species via *P. esculentus* can occur in certain populations (Mezhzherin et al., 2024; Mikulíček et al., 2014; Uzzell et al., 1976). More comprehensive genomic data would be essential to determine if such gene flow is taking place.

Genomic data also would have wider applications. For example, molecular species identification in *Pelophylax* is complicated because *P. esculentus* has no mitochondrial DNA of its own, and mtDNA introgression between the parent species is common in some areas (Plötner et al., 2008). Nuclear markers have the potential to resolve this, but the commonly used marker SAI-1 is not entirely reliable (Dufresnes, Dubey, et al., 2024), and there is a lack of other sequences with known species specific variants.

Target capture sequencing allows for hundreds to thousands of markers to be sequenced in each sample via hybridization with pre-designed RNA baits (Gnirke et al., 2009). As the targeted markers are chosen in advance, it is simple to integrate data from multiple studies. Generally coding DNA and other highly conserved sequences are chosen as markers, resulting in data which is more interesting from a functional perspective and more suitable for comparison between widely divergent lineages (Andermann et al., 2020). Additionally, target capture data can be used to determine the ploidy of individuals (particularly in hybrids) (Weiß et al., 2018). Target capture requires baits to be designed in advance, however as the method is amenable to significantly divergent sequences, it is often feasible to use a bait set designed for one taxon in related taxa or design a bait set covering an extremely large number of species (de Visser et al., 2025; Yardeni et al., 2022). Recently FrogCap, a bait set designed for use across the order Anura, has been successfully used in several studies (Chan et al., 2020; Hutter et al., 2022; Rasolonjatovo et al., 2020). Here we aim to test the usefulness of FrogCap for investigation of the genomics of the taxa comprising the *P. esculentus* hybridogenetic complex.

We gather *Pelophylax* samples from across the Netherlands, including all three taxa. We also include samples from Poland - where the morphology of the taxa appears more distinct (Arntzen, 1981; Berger, 1973). We then use the FrogCap bait set to explore 1) whether (i) it is possible to obtain high coverage genomic data from these samples, 2) (ii) the species can be accurately determined from the molecular data, 3) (iii) the ploidy of the genomes can be identified, and 4) (iv) any nuclear gene flow between *P. lessonae* and *P. ridibundus* can be detected.

## Methods

### Samples

We used samples from a total of 45 individuals in this study, including 38 samples originating from the in the Netherlands (obtained from 19 localities consisting of 18 buccal swabs, 13 skin swabs and 7 tissue samples) and previously reported in Theodoropoulos et al. (2025). Seven samples were obtained from six localities within Poland; these consisted of toe clips or muscle collected with permission from the General Directorate for Environmental Protection (permit nos. DZP-WG.6401.02.2.2018.kp.2 and DZP-WG.6401.75.2023.TŁ.2). The total set of 45 samples included 11 *P. lessonae*, 22 *P. esculentus*, and 12 *P. ridibundus*. Previous barcoding showed that with the exception of one *P. esculentus* all samples from the Netherlands possess *P. lessonae* mtDNA (Theodoropoulos et al., 2025). The seven Polish samples were mitotyped using a PCR-based method that uses *lessonae*- and *ridibundus*-specific primers to amplify fragments of mtDNA that can be verified on an agarose gel (Jośko & Pabijan, 2021). Data for all samples can be found in table S1.

### Library preparation

Concentration of DNA extracts was standardized to a minimum of 100 ng/µl. Library preparation was performed using the NEBNext Ultra™ II FS DNA Library Prep Kit for Illumina (New England Biolabs, MA, USA), using the manufacturer’s instructions with all volumes divided by four. DNA was fragmented enzymatically for 6:30 min before adapter ligation. Size selection targeting a 300 bp insert size was performed using NucleoMag™ beads (Macherey-Nagel, Düren, Germany). Unique combinations of custom i5 and i7 index primers (Integrated DNA Technologies, Leuven, Belgium) were incorporated via nine cycles of PCR. Library concentration and quality were assessed using the Agilent 4200 Tapestation system (Agilent, CA, USA). After library preparation, libraries were pooled equimolarly in batches of 15 with 250 ng of DNA per sample, and vacuum concentrated to 800 ng/µL.

### Target capture sequencing

Target enrichment capture was performed with the FrogCap Ranoidea-v2 marker set (Hutter et al., 2022) (serial code D10260HFRn) using the Mybaits V 5.0 kit (Arbor Biosciences, MI, USA). After a 30 minute pre-hybridisation incubation with the blocking mix, pooled libraries were hybridized at 63 °C for 24 hours. Enriched libraries were bound to streptavidin coated beats and washed to remove non-target DNA before being subject to 14 cycles of PCR using the KAPA HiFi Master Mix (Roche, Basel, Switzerland). 150 bp paired-end sequencing was performed using the NovaSeq 6000 platform (Illumina Inc., San Diego, CA, USA) by BaseClear B.V. (Leiden, Netherlands), targeting 2 Gbp of sequencing data per individual.

### Processing of sequence capture data

We used a bioinformatics pipeline adapted from the NewtCap protocol (de Visser et al., 2025). The upstream data processing was performed via a custom Perl (version v3.38.0) script (*Pipeline_1*.*pl*). Raw demultiplexed reads were trimmed with Trimmomatic (version 0.39) (Bolger et al., 2014) and BBDuk version 38.96 (Bushnell et al., 2017) to remove adapter contamination. BWA-MEM (version 0.7.17) (Li, 2013) was used to map the trimmed reads against reference assembly consisting of the FrogCap Ranoidea-v2 consensus marker sequences (Hutter et al., 2022). The resulting BAM files were processed, deduplicated and genotyped via the *AddOrReplaceReadGroups, MarkDuplicates* and *HaplotypeCaller* functions of GATK (version 4.5.0.0) (McKenna et al., 2010). The resulting VCF files were jointly genotyped with the *GenomicsDBImport* and *GenotypeGVCFs* function of GATK.

Sequencing depth was assessed with a custom R (version 4.4) (R Core Team, 2024) script (*Peakloop2*.*R*) which evaluated the minimum depth of the best covered continuous 100 bp sequence within each target sequence of the reference assembly (this metric was chosen as it is not affected by reference assembly target length). For each individual, the coverage of each marker was collected, and median and total values were calculated. A graph was produced to visualize the performance of each marker in Rstudio (version 24.04.2) (Posit team, 2025) using the packages ggplot2, patchwork and tidyr (Pedersen, 2025; Wickham, 2016; Wickham et al., 2025) The median coverage of each marker was calculated and markers were then ordered by the sum of their median coverage.

The joint VCF file was filtered using VCFtools (version 0.1.16) (Danecek et al., 2011) to remove indels, sites with a minor allele frequency of less than 0.1 and sites with missing data, sequencing depth of less than 10, or genotype quality of less than 20, in any individual. Finally the *–thin* function was used to select one SNP per reference sequence.

### Principal component analysis

Principal Component Analyses (PCAs) were performed using the R packages gdsfmt and SNPRelate (Zheng et al., 2012). An initial PCA was performed on the entire dataset (all 45 samples) and then separate PCAs were performed on each of the three taxa targeted in this study (11 *P. lessonae*, 22 *P. esculentus*, and 12 *P. ridibundus*). The results of the analysis were visualized with the ggplot2 package (Wickham, 2016).

### Admixture analysis

ADMIXTURE (Alexander et al., 2009) was used to determine the ancestry of each sample. Input consisted of a VCF file containing one random SNP per marker, filtered for sites with no missing data. As *P. esculentus* is a hybrid of the two other taxa involved in this study, the number of ancestral gene pools (K) was set to two (as there are only two nuclear lineages of interest within the study population). Twenty-five replicates of the analysis were combined with the program CLUMPAK (Kopelman et al., 2015) and the results were visualized with the ggplot2 package (Wickham, 2016).

### Heterozygosity/ancestry analysis

After importing VCF data with the vcfR package (Knaus & Grünwald, 2017), the triangulaR (Wiens et al., 2025) package was used to calculate the hybridity and heterozygosity based on SNPs that were 100% diagnostic between the parental species (i.e. SNPs that were present in all 11 *P. lessonae* individuals and absent in all 12 *P. ridibundus* individuals, or vice versa). The data was then visualized with the ggplot2 package (Wickham, 2016).

### Ploidy analysis

The species-specific SNPs used to determine hybridity were also employed to determine the ploidy and parental contribution of the *P. esculentus* samples. For each SNP locus in each sample the ratio of the sequencing depth of the *P. lessonae* allele to the total sequencing depth was calculated. For diploid *P. esculentus* samples these ratios should average 0.5, for triploid the ratios should be either 0.33 or 0.66 depending on whether the individual possesses an extra copy of the *P. ridibundus* or *P. lessonae* or genome. For each sample the frequency of SNPs was plotted against the calculated allele ratio.

Additionally, the ploidy of all samples was assessed using the program nQuire (cloned from Git commit 8975c94) (Weiß et al., 2018). The BAM files produced for each sample were used as input to create .bin files via the *create* function with -c set to 20 and -p to 20. The *denoise* function was then used on the .bin files, followed by ploidy model fitting with the *histotest* and *view* functions. The resulting allele ratio distributions were normalized and combined. Gaussian mixture models were utilized to assess the ploidy level, where a free model is compared to models fixed on diploidy, triploidy and tetraploidy. The model with the lowest difference from the free model was interpreted as the most likely ploidy of the sample. The results were visualized with the ggplot2 package (Wickham, 2016).

## Results

### Sequence capture performance

We obtained 134.9 Gbp of raw sequence data, with a mean of 11,074,445 (s.d. 5,009,066) read pairs per sample. A mean of 69.4% (s.d. 8.6%) are duplicates and 37.2% (s.d. 15.2%) of reads can be mapped to the target sequences, resulting in a mean of 11.4% of read pairs being useful (1,260,626 read pairs per sample). After deduplication and mapping median peak coverage varies between samples from a minimum of 9 to a maximum 81 (mean 35.4, s.d. 23.4).

Coverage varies between target sequences (Data S1). Across all samples, 2395 of 12909 markers (18%) have a median coverage across all samples of over 100, and 4407 markers (34%) have less than 10. VCF filtering resulted in a final selection of 1957 SNPs available for further analysis.

### Principal component analysis

In the PCA including all 45 samples (Fig. 1A), three distinct groups are separated by the first principal component (PC1), which explains 33% of the total variation in the dataset. PC1 corresponds to *lessonae*-*ridibundus* variation, with all *P. esculentus* samples aligning along the origin of this axis, and all *P. lessonae* and *P. ridibundus* located in tight clusters equidistant from the origin. A single *P. lessonae* individual (swab_1673) is the only sample to vary significantly along the PC2 axis (which accounts for 7.5% of total variation). To reveal any variation obscured by this sample, we repeat the PCA with it excluded (Fig. 1B), but this does not show a significant effect on the overall clustering of samples. When separate PCAs (Fig. 1C-F), are performed for each taxon, we observe that the Polish individuals form a distinct cluster in all three cases. We also observe a clustering of triploid vs. diploid samples among Dutch *P. esculentus* (see the ploidy analysis below).

**Fig 1.**
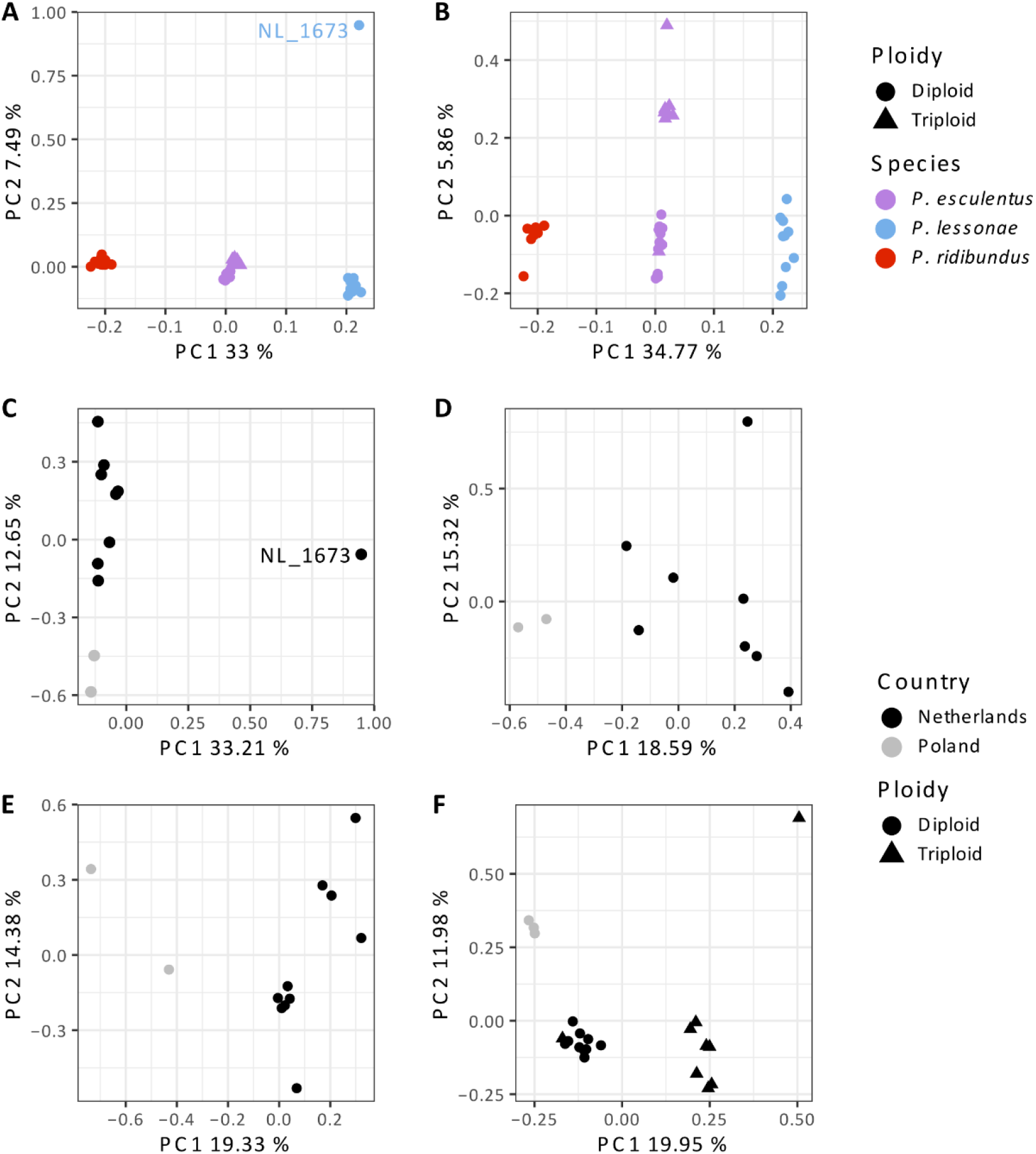
Principal Component Analysis of waterfrogs (*Pelophylax*). **A)** PCA including all samples used in this study, the three taxa are separated by the first principal component. A single *P. lessonae* (NL_1673) accounts for the major variation along the second principal component, with all other samples tightly clustered by taxa. **B)** PCA including all samples other than NL_1673, the first principal component is almost unchanged, but the taxa diverge along the second principal component. **C)** All *P. lessonae* samples. **D)** *P. lessonae* excluding NL_1673. **E)** all *P. ridibundus* samples. **F)** All *P. esculentus* samples.

### Admixture analysis

All *P. ridibundus* and *P. lessonae* individuals have complete *P. ridibundus* and *P. lessonae* ancestry, respectively (Fig. 2). All *P. esculentus* samples show a mixed ancestry, however in several the contribution from *P. lessonae* is slightly greater than the expected 50%. This is particularly the case in triploid samples that carry two copies of the *P. lessonae* genome.

**Fig 2.**
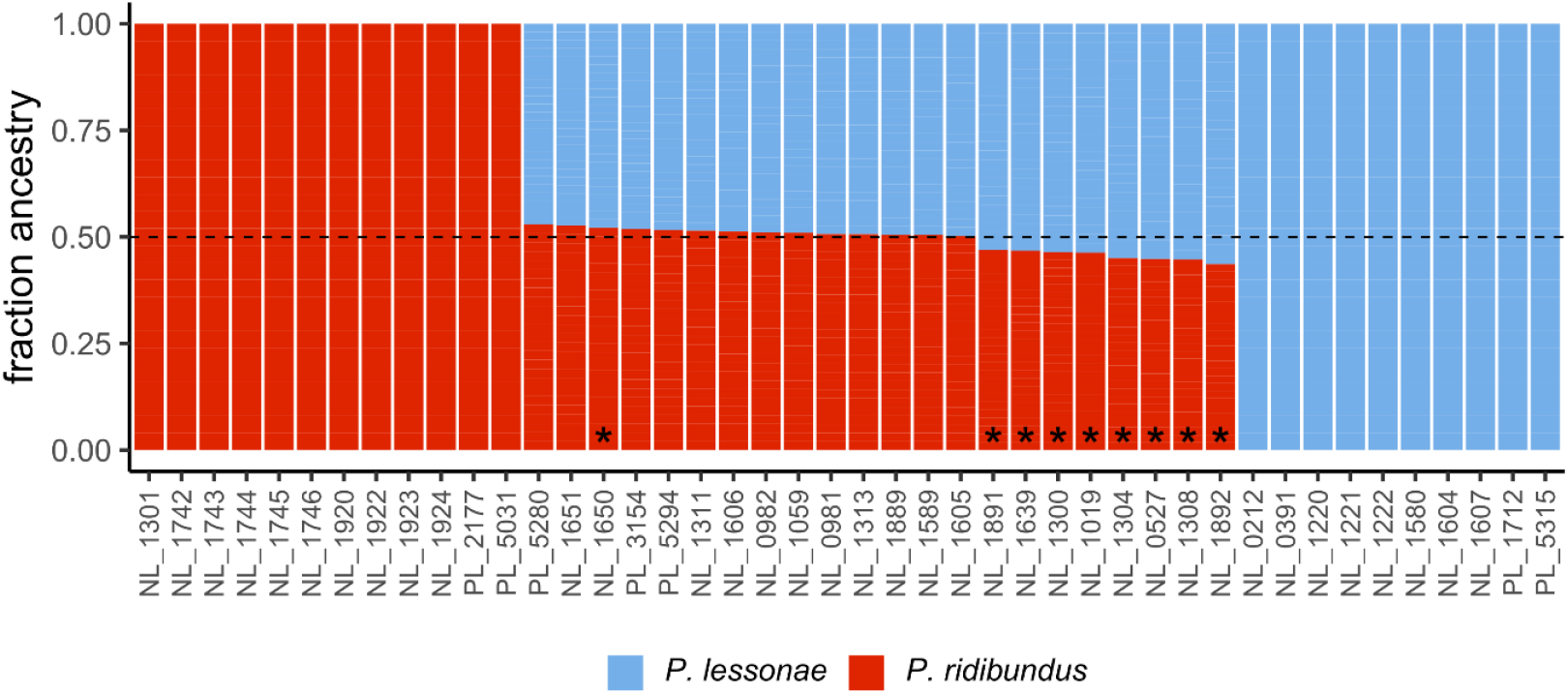
Admixture analysis of waterfrogs (*Pelophylax*). Triploid samples are denoted with an asterisk (**\***). All samples have either 100% ancestry from one of the parent species, or close to 50% ancestry from both. The triploid *P. esculentus* samples that show slightly higher *P. lessonae* contribution all have two copies of the *P. lessonae* genome, whereas the remaining triploid has two copies of the *P. ridibundus* genome.

### Ancestry/heterozygosity analysis

696 SNPs were species diagnostic in all individuals from the parental species. All *P. esculentus* samples cluster in the upper vertex of the triangle plot (Fig. 3) – with a hybrid index (ancestry) close to 0.50 and near 100% interspecies heterozygosity. This is the expected behaviour of F1 hybrids, which *P. esculentus* effectively are. However, corresponding to the admixture analysis some *P. esculentus*, particularly triploids with two copies of the *P. lessonae* genome, are slightly displaced in the direction of *P. lessonae*, with swab_1304 and swab_1892, the triploids with the lowest coverage, showing the greatest displacement.

**Fig 3.**
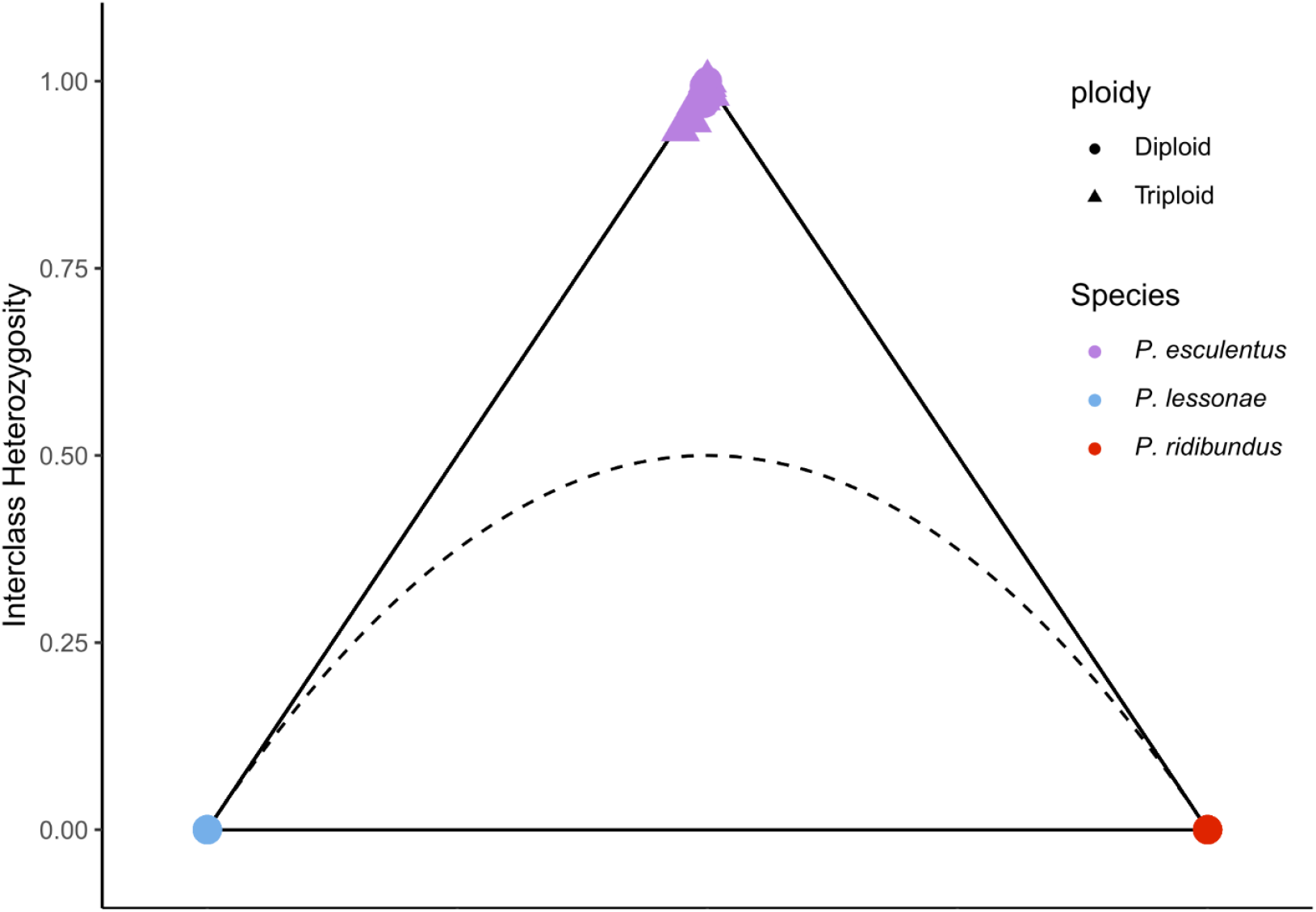
Heterozygosity/hybridity analysis of waterfrogs (*Pelophylax*). The hybrid index (x-axis) indicates what percentage of the markers of an individual belong to either parent species (with markers with alleles from both parental species being scored as 0.5, regardless of the actual allele ratio), where 0 represents *P. lessonae* and 1 *P. ridibundus*. The y-axis indicates the interclass heterozygosity, meaning the proportion of loci with alleles from both parental groups. The dotted black curve indicates the boundary below which individuals cannot occur, assuming Hardy-Weinberg Equilibrium. All *P. esculentus* samples behave similarly to an ideal F1 hybrid (with all markers being heterozygous and having 50% ancestry from each parental species).

### Ploidy analysis

We observe that 13 of the 22 *P. esculentus* samples show allele ratios normally distributed around 0.5 for the SNPs diagnostic for the parent species, typical of diploidy (Fig. 4). Of the nine remaining *P. esculentus* samples, eight show allele ratios of 2:1 in favour of *P. lessonae*, indicating that they are triploids with two copies of the *P. lessonae* genome. The remaining sample is also triploid but with allele ratios 2:1 in favour of *P. ridibundus*.

**Fig 4.**
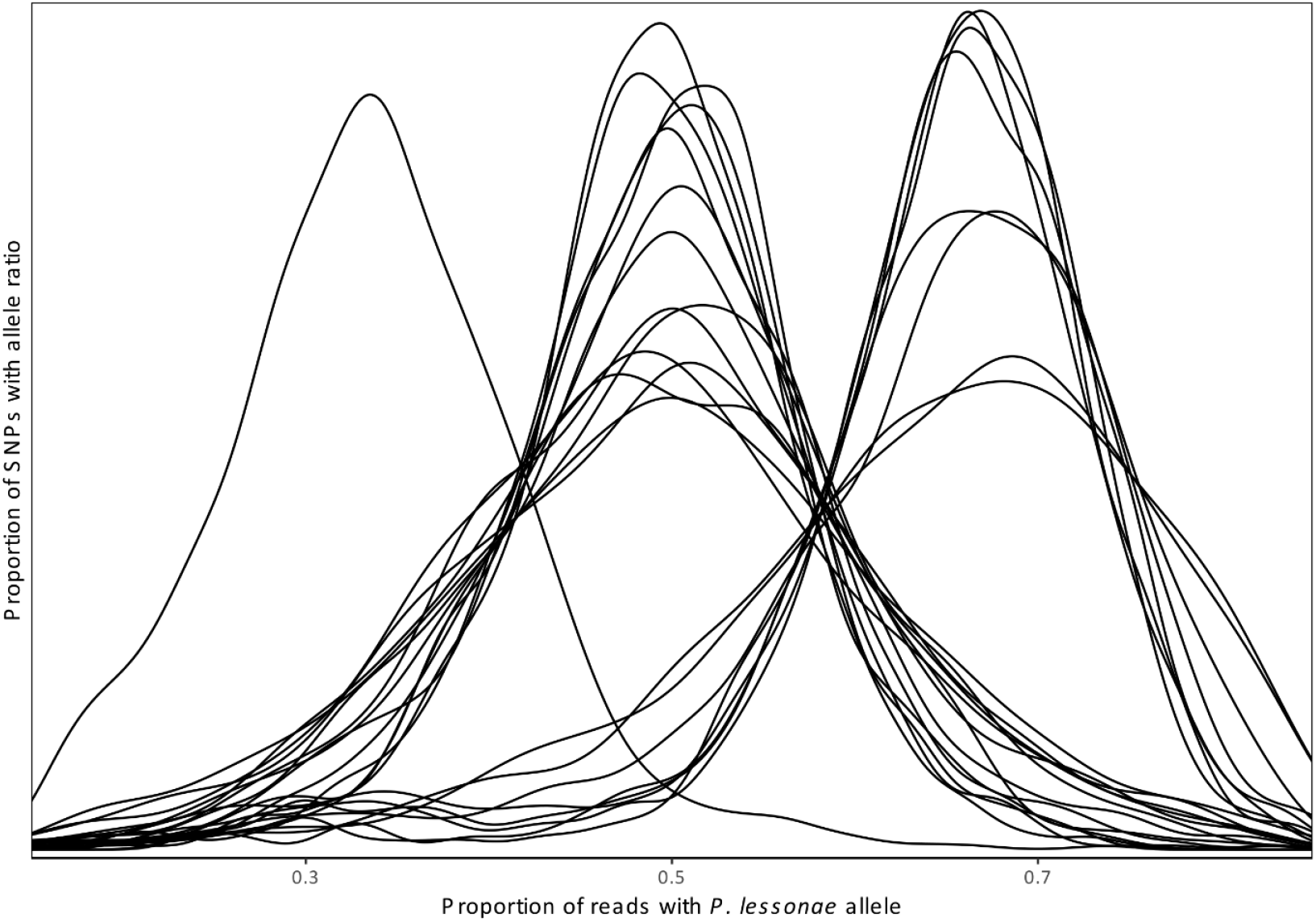
Polyploidy and genome composition of *Pelophylax esculentus* samples. For diploid samples the proportion of reads with *P. lessonae* specific SNPs is normally distributed around 0.5. Eight samples instead show a mean ratio of approximately 0.66, indicating that they are triploid with two copies of the *P. lessonae* genome and a single copy of the *P. ridibundus* genome. A single sample has the opposite genome composition, as indicated by its peak at approximately 0.33.

Analysis of the *P. esculentus* data with nQuire confirms the samples consist of nine triploids and 13 diploids. All *P. lessonae* and *P. ridibundus* samples are diploid (Fig. S1-3). The number of markers available for ploidy analyses varies with both overall coverage and heterozygosity, resulting in lower *R*^*2*^ values for the parental species (which are less heterozygous than the parents). However, while the *P. lessonae* and *P. ridibundus* samples typically have fewer than half the informative markers compared to *P. esculentus* samples, there is enough data available for confident ploidy determination in all samples.

### Distribution within the Netherlands

We find *P. ridibundus* samples only within the western part of the Netherlands (Fig. 5), consistent with their described natural range. The majority of *P. lessonae* samples originate from the eastern Netherlands however we find that two localities within the southern part of the Dutch coastal dune areas are also populated by *P. lessonae*. We observe the presence of *P. esculentus* throughout the Netherlands, however all samples in the west are triploids with two copies of the *P. lessonae* genome, in the east all samples are diploid apart from the single triploid with two copies of the *P. ridibundus* genome.

**Fig 5.**
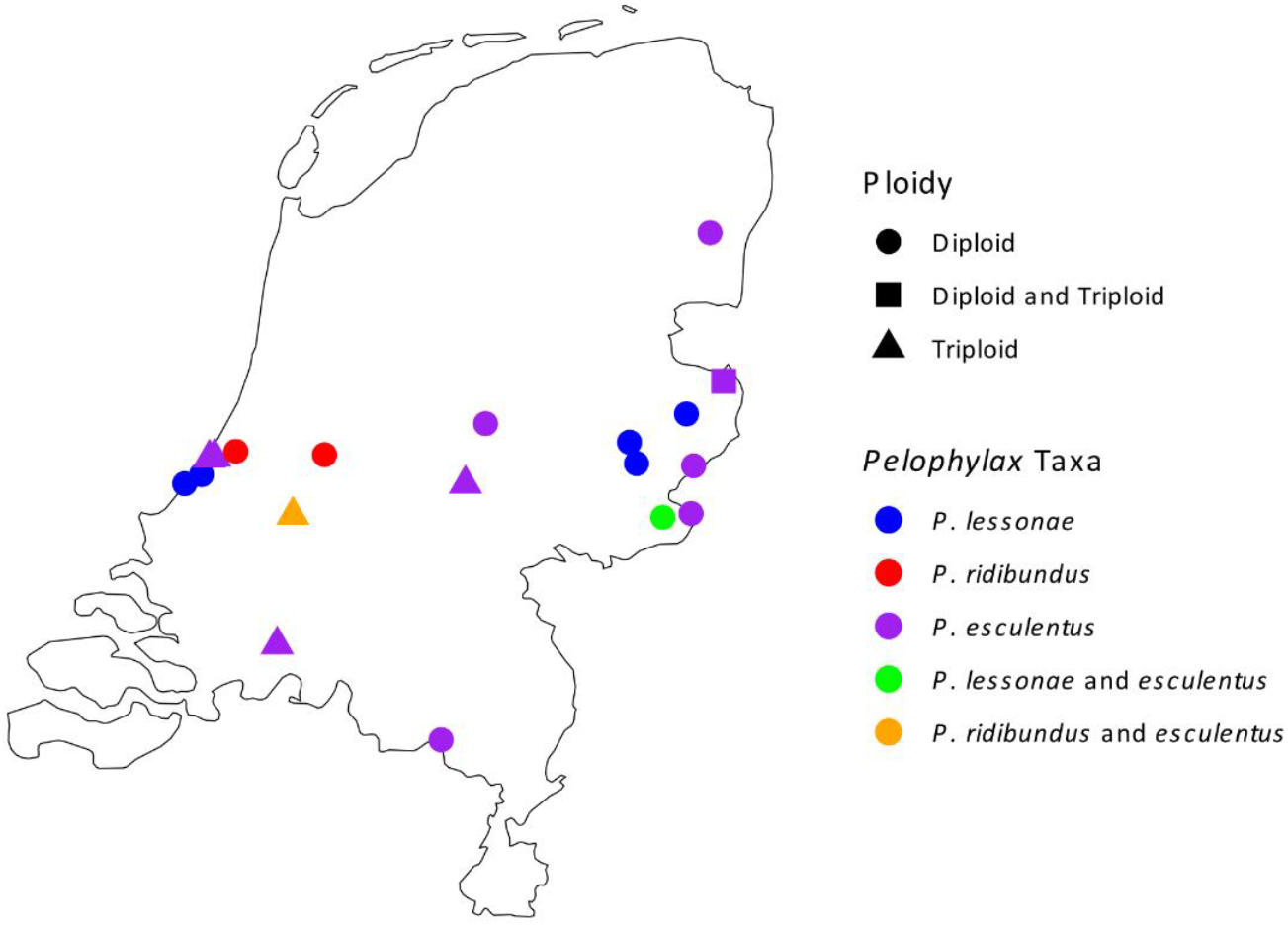
Map of waterfrog (*Pelophylax*) localities sampled in the Netherlands. *P. esculentus* occurs throughout the country, *P. ridibundus* is found in its natural range in the west. Two localities in the coastal dune areas are populated with *P. lessonae*, outside of its typical distribution in the east. All *P. esculentus* found in western areas are triploid with two copies of the *P. lessonae* genome. The sole triploid from the east instead has two copies of the *P. ridibundus* genome.

## Discussion

In this study we examine whether the target capture bait set FrogCap could be effectively used to interrogate the *Pelophylax esculentus* hybridogenetic system, including questions regarding sample identification, ancestry, introgression and polyploidy.

The bait set used within this study (FrogCap Ranoidea v2) (Hutter et al., 2022) targets 12,909 markers, which are highly heterogeneous in terms of length (from 100 to 12,000 bp) and include exons, introns and various non-coding ultra conserved elements. While based on transcripts from species across the Ranoidea superfamily, no sequences from *Pelophylax* were used in the bait design. Therefore, it is unsurprising that 34% of markers perform poorly in *Pelophylax*. Nevertheless, in our dataset, over 8,500 marker sequences have a region of at least 100 bp with at least 10x coverage in at least 50% of individuals, sufficient for reliable phylogenetic and phylogeographic analyses. We also note that the authors of Frogcap designed an additional bait set, Reduced-Ranoidea (Hutter et al., 2022), that targets 3,247 markers, which are a subset of the full Ranoidea set. Given that a smaller bait set is less costly and is likely to result in higher target coverage per Gbp of sequence data produced (as approximately the same number of captured reads will be spread across fewer target bases), this may also be useful for investigations into *Pelophylax*.

We show that target capture data allows for very clear differentiation of the taxa involved in the *P. esculentus* hybridogenetic complex. This is useful in its own right, as identification through morphology or vocalization is often difficult, especially in larvae and juveniles (Arntzen, 1981; Meilink et al., 2024). Additionally, previous molecular species identification methods have proven unreliable (Dufresnes, Dubey, et al., 2024). While target capture sequencing is more resource intensive than might be desired for simple species identification, the resulting data allows for identification of a large number of species-specific polymorphisms, which can then be used to design PCR-based protocols such as KASP genotyping (Semagn et al., 2014) that may prove more reliable than those currently available (Dufresnes, Dubey, et al., 2024).

We observe that polyploidy is easily recognizable using the FrogCap marker set. Compared to cytogenetic techniques or flow cytometry this provides a simple method to determine the ploidy of a large number of samples. Additionally, we show that the same data can also show which of the parental species is present in two copies in each sample. The identification of reliable species-specific SNPs also creates the possibility of using techniques such as droplet digital PCR (Hindson et al., 2011) to identify triploids and the genome composition on a large scale. However, we note that ploidy determination in our samples was based only on allele depth ratios. While the analysis appears robust in this exploratory study, it may still be useful to verify this methodology by genotyping individuals of known ploidy assessed by independent cytogenetic or flow-cytometry analyses.

Our data grant insight into the distribution of *Pelophylax* taxa within the Netherlands. Overall species identification matches the distribution described in the literature (Creemers & van Delft, 2009). All *P. ridibundus* samples originated from localities within the west of the Netherlands, within their natural range. While most *P. lessonae* were found in the east of the Netherlands, we confirm previous observations of populations in the coastal dune area, approximately 50 km east of their traditionally recognized native range, and 70 km south of a previously recognized introduction in North Holland (NDFF, 2015). Several non-native species of amphibians are known to have been introduced to the dune area (Koster et al., 2022; Kuijt et al., 2022; Robbemont et al., 2023; Vliegenthart et al., 2022), and it is therefore likely that *P. lessonae* represents a further example. In agreement with previous studies employing cytogenetic data (Blommers-Schlösser, 1990), we observe a marked pattern in the ploidy of *P. esculentus* samples, in the west, co-occurring with *P. ridibundus*, all samples were triploids with two copies of the *P. lessonae* genome. In the east, inside the natural range of *P. lessonae*, most *P. esculentus* were diploid, with only a single triploid observed (and this with the opposite configuration – two copies of the ridibundus genome). It is worth noting that our study does not include samples from the northern and central part of the range of *P. ridibundus* within the Netherlands, which would be useful to include in any future research into the population genetics of *Pelophylax*.

In a previous study (Theodoropoulos et al., 2025) we observed that *P. lessonae* mtDNA was extremely widespread throughout *Pelophylax* samples throughout the Netherlands regardless of which *Pelophylax* taxon the sample was assigned to. Our results in this study confirm that this is due to mtDNA introgression of *P. lessonae* mtDNA into *P. ridibundus* via *P. esculentus*. The result is that, except for a single *P. esculentus*, all individuals, including all *P. ridibundus*, possess *P. lessonae* mtDNA. Unidirectional introgression of mtDNA is the expected result of the *lessonae*-eliminating system present in the Netherlands. As *P. esculentus* typically mate with *P. lessonae* generation after generation, it is highly likely that they will eventually inherit *P. lessonae* mtDNA. When a *P. esculentus* female carrying *P. lessonae* mtDNA mates with *P. ridibundus*, the result will be an individual with purely *P. ridibundus* nuclear DNA but *P. lessonae* mtDNA (Plötner et al., 2008).

We also found that several *P. esculentus* samples appear to have slightly greater than 50% *P. lessonae* ancestry. This could potentially be seen as evidence of nuclear gene flow (Mezhzherin et al., 2024; Mikulíček et al., 2014; Uzzell et al., 1976), especially as the mechanics of the hybridogenetic system mean that any introgression of nuclear DNA is likely to follow the same pattern as mtDNA, against the direction of genome elimination. However, we believe this is more likely to be an artifact due to miscalled haplotypes. The affected samples are all triploids with two copies of the *P. lessonae* genome, and the two samples with the largest excess *P. lessonae* contribution are the two triploids with the lowest overall coverage, and therefore the individuals most likely to be affected by miscalled haplotypes where the *P. ridibundus* allele is not recorded.

While *P. ridibundus* and *P. lessonae* are the only ancestral lineages natively present within the Netherlands (and hence examined in this study), the genus *Pelophylax* includes other taxa capable of forming hybridogenetic complexes, for example the Iberian frog *P. grafi*, a hybridogenetic hybrid between *P. perezi* and *P. ridibundus* (Graf et al., 1977). The interactions between these taxa are less well studied, especially when their native ranges do not border each other. However, due to the large number of anthropogenic *Pelophylax* introductions in recent times (Denoël & Dufresnes, 2025; Doniol-Valcroze et al., 2021; Dufresnes & Dubey, 2020; Papežík et al., 2024) these exotic systems may become extremely important. The ability to exclude a competing genome from gametogenesis could allow invasive lineages to spread through, and displace, native populations extremely rapidly. Consequently, *Pelophylax* has been identified as one of the taxa most vulnerable to disruption by introduced species, creating a critical need for monitoring (Dufresnes & Mazepa, 2020).

Unfortunately, it is not always easy to determine whether an introduction has taken place, given the plastic morphology of *Pelophylax*, and limitations of current molecular approaches. The target capture approach we test in this study offers a powerful tool for examining the ancestral contribution of many lineages to any given population. A possible example of interest is the sample swab_1673, which appears highly divergent from other *P. lessonae* samples in the PCA. As this divergence was orthogonal to the otherwise dominant *P. lessonae* - *P. ridibundus* variation, it cannot easily be explained by interaction between the native taxa. However, this divergence may represent introgression from a non-native *Pelophylax* taxon, which could be confirmed if the target capture methodology was applied to a wider selection of *Pelophylax* taxa.

We conclude that target capture sequencing employing the FrogCap marker set (Hutter et al., 2022) is an effective method of gathering large-scale genetic information from samples within the genus *Pelophylax*. We recommend its use for future studies investigating the hybridogenetic complexes and other evolutionary and ecological questions in the genus.

## Data availability

All raw reads can be found as a part of the NCBI accession associated with Bioproject: PRJNA1330369. All scripts used for the processing of data in this study are available at the projects GitHub repository: https://github.com/Wielstra-Lab/Pelophylax_target_capture.

## Acknowledgements

We give thanks to the volunteers and staff of Reptile, Amphibian and Fish Conservation Netherlands (RAVON) for their support of this research. This work was performed using the ALICE compute resources provided by Leiden University.

