## Supplementary figures and tables for "Target capture sequencing provides insights into hybridogenetic water frogs"

**Table of Contents:**

| | |
|---|---|
| **Figure S1: Ploidy of *Pelophylax ridibundus* samples** | Page 2 |
| **Figure S2: Ploidy of *Pelophylax lessonae* samples** | Page 3 |
| **Figure S3: Ploidy of *Pelophylax esculentus* samples** | Page 4 |
| **Table S1: Sample details** | Page 5-6 |

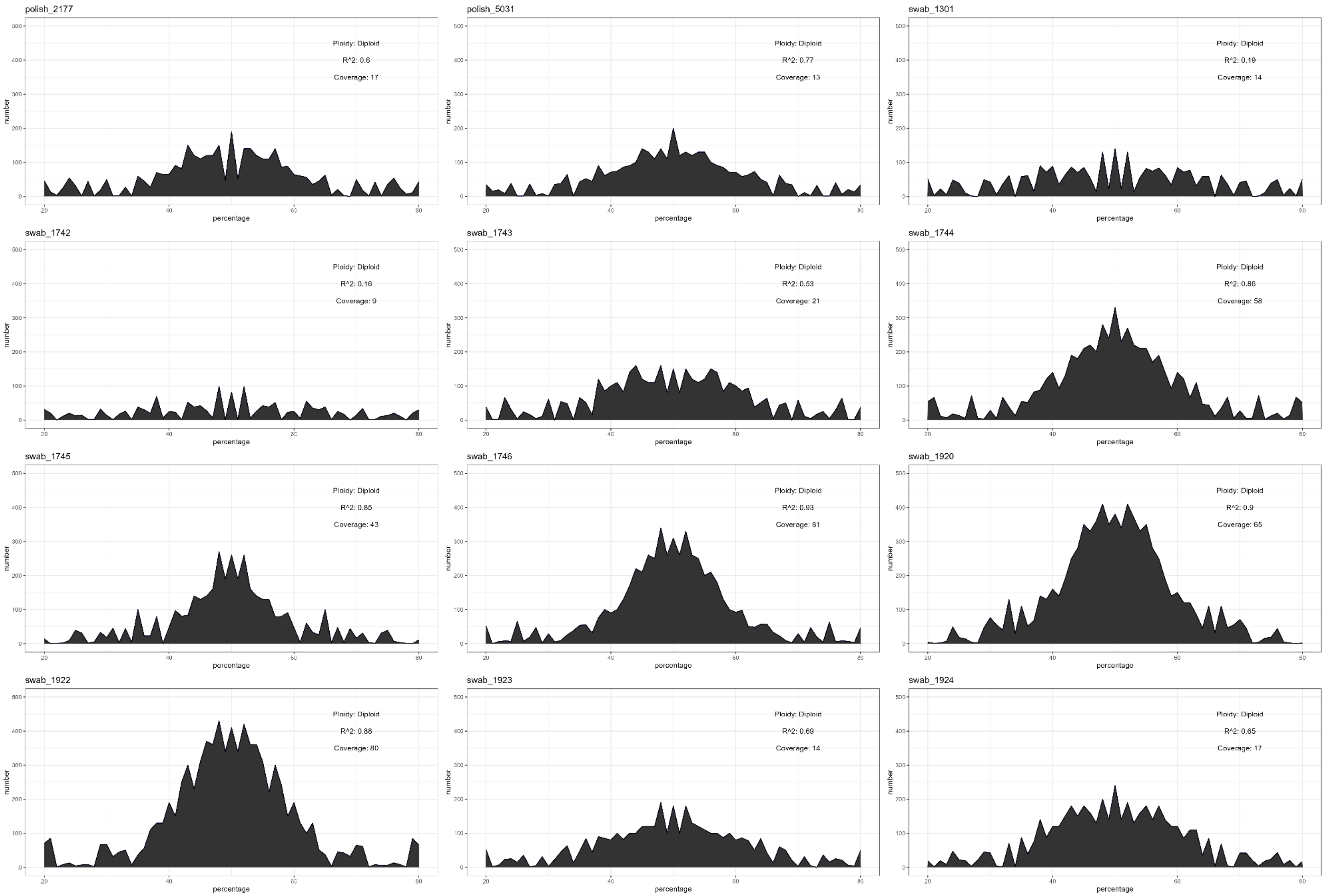

**Fig S1. Ploidy of *Pelophylax ridibundus* samples**. In each subfigure, the most likely model of ploidy is noted, along with the median coverage (over all markers) and the $R^2$ value for the best ploidy model. The x-axis displays the allele ratio of each SNP. The y-axis shows number of SNPs with a given allele ratio. All *P. ridibundus* samples were found to be diploid

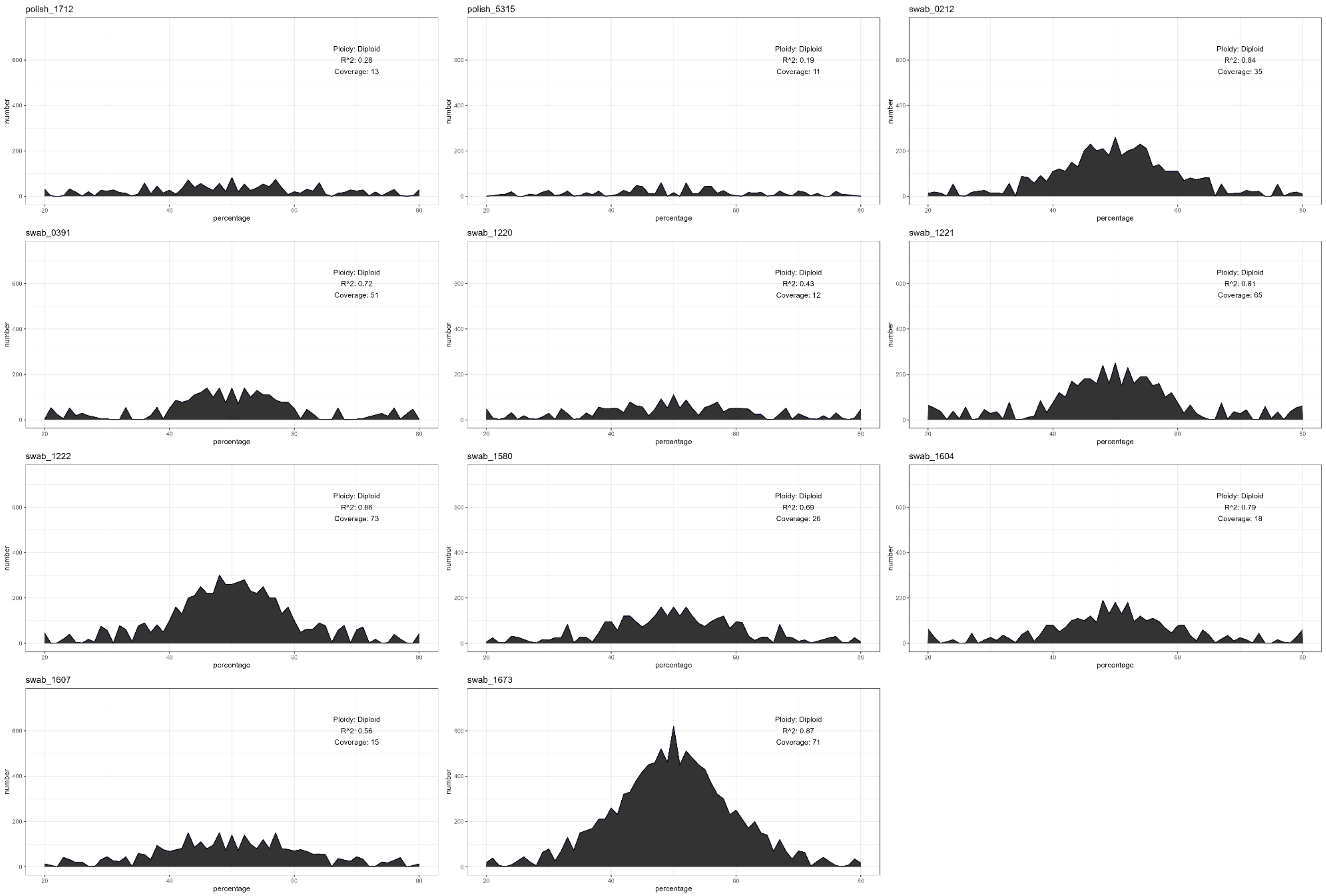

**Fig S2. Ploidy of *Pelophylax lessonae* samples**. In each subfigure, the most likely model of ploidy is noted, along with the median coverage (over all markers) and the $R^2$ value for the best ploidy model. The x-axis displays the allele ratio of each SNP. The y-axis shows number of SNPs with a given allele ratio. All *P. lessonae* samples were found to be diploid

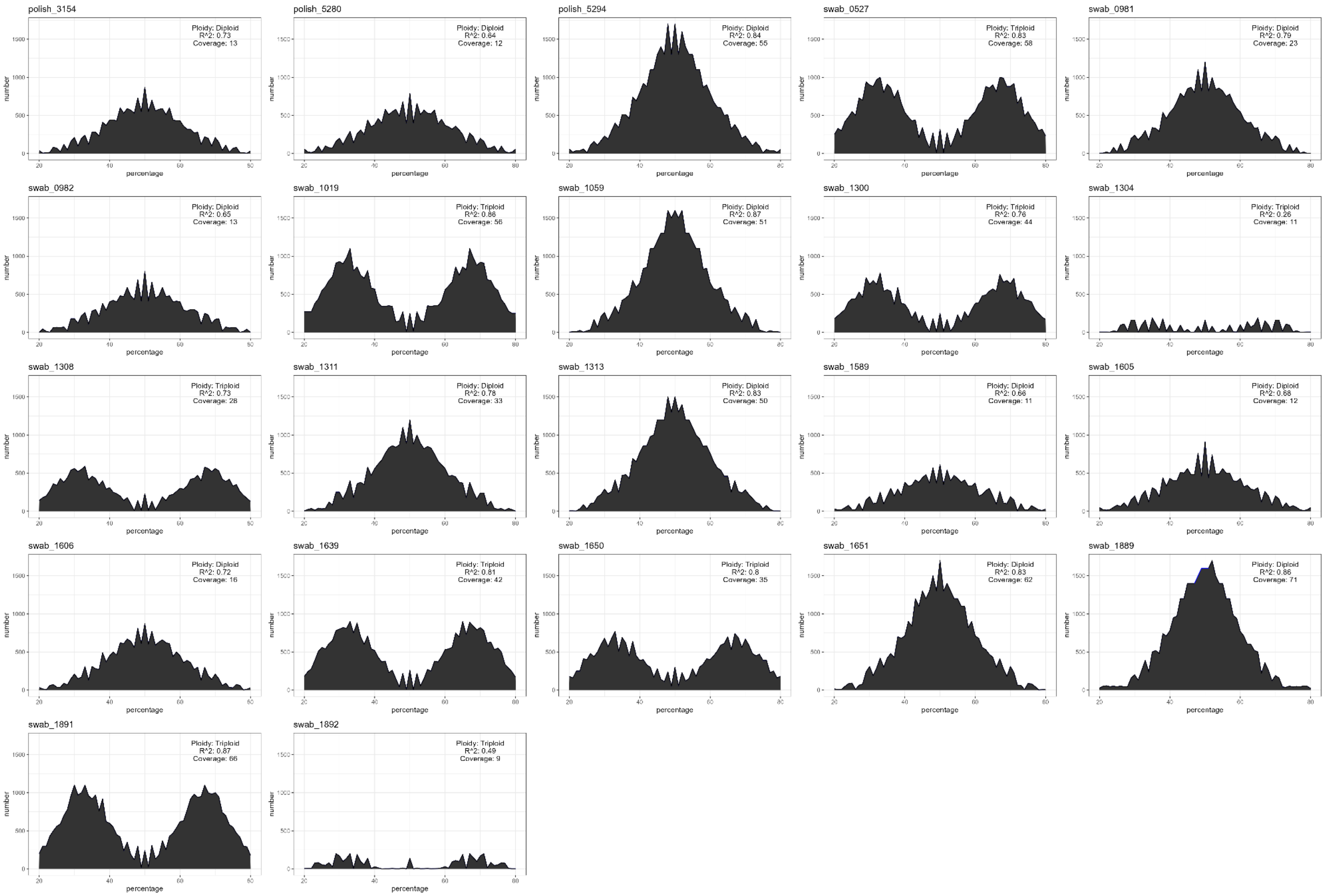

**Fig S3. Ploidy of *Pelophylax esculentus* samples**. In each subfigure, the most likely model of ploidy is noted, along with the median coverage (over all markers) and the $R^2$ value for the best ploidy model. The x-axis displays the allele ratio of each SNP. The y-axis shows number of SNPs with a given allele ratio. 9 *P. esculentus* samples were found to be triploid and the rest diploid.

**Table S1. Details of samples used in this study**. Note that all samples from the Netherlands are also reported in (Theodoropoulos et al., 2025).

| Sample ID | Alternative ID | Sample type | Country | Locality | Latitude | Longitude | mtDNA | Taxon (PCA) | Ploidy | Copy number | | SRA accession number |
|---|---|---|---|---|---|---|---|---|---|---|---|---|
| | | | | | | | | | | *P. lessonae* | *P. ridibundus* | |
| PL_1712 | MPFC1712 | Tissue | Poland | Klewiny | 54.2834 | 22.0877 | *P. lessonae* | *P. lessonae* | Diploid | 2 | 0 | |
| PL_5315 | MPFC5315 | Tissue | Poland | Straż | 53.3395 | 23.3711 | - | *P. lessonae* | Diploid | 2 | 0 | |
| PL_5031 | MPFC5031 | Tissue | Poland | Mydlniki | 50.0843 | 19.8403 | *P.ridibundus* | *P.ridibundus* | Diploid | 0 | 2 | |
| PL_2177 | MPFC2177 | Tissue | Poland | Stawy Bugaj | 49.9813 | 19.4250 | *P.ridibundus* | *P.ridibundus* | Diploid | 0 | 2 | |
| PL_3154 | MPFC3154 | Tissue | Poland | Przeręb | 50.0145 | 19.4056 | *P.esculentus* | *P.esculentus* | Diploid | 1 | 1 | |
| PL_5280 | MPFC5280 | Tissue | Poland | Przeręb | 50.0145 | 19.4056 | *P.esculentus* | *P.esculentus* | Diploid | 1 | 1 | |
| PL_5294 | MPFC5294 | Tissue | Poland | Ispina | 50.1132 | 20.3581 | *P.esculentus* | *P.esculentus* | Diploid | 1 | 1 | |
| NL_1300 | swab_1300 | Buccal swab | Netherlands | Bilwijk Krimpenerwaard | 51.9871 | 4.7836 | *P. lessonae* | *P.esculentus* | Triploid | 2 | 1 | |
| NL_1301 | swab_1301 | Buccal swab | Netherlands | Bilwijk Krimpenerwaard | 51.9871 | 4.7836 | *P. lessonae* | *P. ridibundus* | Diploid | 0 | 2 | |
| NL_1673 | swab_1673 | Skin swab | Netherlands | Borne | 52.2986 | 6.7625 | *P. lessonae* | *P. lessonae* | Diploid | 2 | 0 | |
| NL_0391 | swab_0391 | Skin swab | Netherlands | Den Haag, Westduinpark | 52.0833 | 4.2393 | *P. lessonae* | *P. lessonae* | Diploid | 2 | 0 | |
| NL_1220 | swab_1220 | Skin swab | Netherlands | Den Haag, Westduinpark | 52.0833 | 4.2393 | *P. lessonae* | *P. lessonae* | Diploid | 2 | 0 | |
| NL_1221 | swab_1221 | Skin swab | Netherlands | Den Haag, Westduinpark | 52.0833 | 4.2393 | *P. lessonae* | *P. lessonae* | Diploid | 2 | 0 | |
| NL_1222 | swab_1222 | Skin swab | Netherlands | Den Haag, Westduinpark | 52.0833 | 4.2393 | *P. lessonae* | *P. lessonae* | Diploid | 2 | 0 | |
| NL_1019 | swab_1019 | Buccal swab | Netherlands | Duinoord | 52.1630 | 4.3637 | *P. lessonae* | *P.esculentus* | Triploid | 2 | 1 | |
| NL_1311 | swab_1311 | Buccal swab | Netherlands | Gastelsche Heide | 51.2930 | 5.5289 | *P. lessonae* | *P.esculentus* | Diploid | 1 | 1 | |
| NL_1313 | swab_1313 | Buccal swab | Netherlands | Gastelsche Heide | 51.2930 | 5.5289 | *P. lessonae* | *P.esculentus* | Diploid | 1 | 1 | |
| NL_1059 | swab_1059 | Tissue | Netherlands | Haaksbergerveen | 52.1387 | 6.7984 | *P. lessonae* | *P.esculentus* | Diploid | 1 | 1 | |
| NL_1605 | swab_1605 | Tissue | Netherlands | Haaksbergerveen | 52.1196 | 6.7837 | *P. lessonae* | *P.esculentus* | Diploid | 1 | 1 | |
| NL_1889 | swab_1889 | Buccal swab | Netherlands | Landgoed Staverden | 52.2688 | 5.7526 | *P. lessonae* | *P.esculentus* | Diploid | 1 | 1 | |
| NL_1304 | swab_1304 | Buccal swab | Netherlands | Lentevreugd | 52.1658 | 4.3900 | *P. lessonae* | *P.esculentus* | Triploid | 2 | 1 | |
| NL_1308 | swab_1308 | Buccal swab | Netherlands | Lentevreugd | 52.1658 | 4.3900 | *P. lessonae* | *P.esculentus* | Triploid | 2 | 1 | |
| NL_0527 | swab_0527 | Buccal swab | Netherlands | Liesbos | 51.5866 | 4.7047 | *P. ridibundus* | *P.esculentus* | Triploid | 2 | 1 | |

| | | | | | | | | | | | |
|---|---|---|---|---|---|---|---|---|---|---|---|
| NL_1639 | swab_1639 | Buccal swab | Netherlands | Lunterse Buurtbos | 52.0832 | 5.6513 | *P. lessonae* | *P.esculentus* | Triploid | 2 | 1 |
| NL_1891 | swab_1891 | Buccal swab | Netherlands | Lunterse Buurtbos | 52.0867 | 5.6513 | *P. lessonae* | *P.esculentus* | Triploid | 2 | 1 |
| NL_1892 | swab_1892 | Buccal swab | Netherlands | Lunterse Buurtbos | 52.0867 | 5.6513 | *P. lessonae* | *P.esculentus* | Triploid | 2 | 1 |
| NL_1589 | swab_1589 | Skin swab | Netherlands | Masterveld | 51.9918 | 6.7858 | *P. lessonae* | *P.esculentus* | Diploid | 1 | 1 |
| NL_0212 | swab_0212 | Buccal swab | Netherlands | Meijendel TNO | 52.1114 | 4.3242 | *P. lessonae* | *P. lessonae* | Diploid | 2 | 0 |
| NL_1650 | swab_1650 | Skin swab | Netherlands | Plooi Tilligte | 52.4000 | 6.9500 | *P. lessonae* | *P.esculentus* | Triploid | 1 | 2 |
| NL_1651 | swab_1651 | Skin swab | Netherlands | Plooi Tilligte | 52.4000 | 6.9500 | *P. lessonae* | *P.esculentus* | Diploid | 1 | 1 |
| NL_1920 | swab_1920 | Buccal swab | Netherlands | Poelgeest | 52.1832 | 4.4972 | *P. ridibundus* | *P. ridibundus* | Diploid | 0 | 2 |
| NL_1922 | swab_1922 | Buccal swab | Netherlands | Poelgeest | 52.1832 | 4.4972 | *P. ridibundus* | *P. ridibundus* | Diploid | 0 | 2 |
| NL_1923 | swab_1923 | Tissue | Netherlands | Poelgeest | 52.1832 | 4.4972 | *P. ridibundus* | *P. ridibundus* | Diploid | 0 | 2 |
| NL_1924 | swab_1924 | Buccal swab | Netherlands | Poelgeest | 52.1832 | 4.4972 | *P. ridibundus* | *P. ridibundus* | Diploid | 0 | 2 |
| NL_1742 | swab_1742 | Skin swab | Netherlands | Kockengen | 52.1728 | 4.9425 | *P. lessonae* | *P. ridibundus* | Diploid | 0 | 2 |
| NL_1743 | swab_1743 | Skin swab | Netherlands | Kockengen | 52.1728 | 4.9425 | *P. lessonae* | *P. ridibundus* | Diploid | 0 | 2 |
| NL_1744 | swab_1744 | Skin swab | Netherlands | Kockengen | 52.1728 | 4.9425 | *P. lessonae* | *P. ridibundus* | Diploid | 0 | 2 |
| NL_1745 | swab_1745 | Skin swab | Netherlands | Kockengen | 52.1728 | 4.9425 | *P. lessonae* | *P. ridibundus* | Diploid | 0 | 2 |
| NL_1746 | swab_1746 | Skin swab | Netherlands | Kockengen | 52.1728 | 4.9425 | *P. lessonae* | *P. ridibundus* | Diploid | 0 | 2 |
| NL_0981 | swab_0981 | Buccal swab | Netherlands | Valthe | 52.8571 | 6.8809 | *P. lessonae* | *P.esculentus* | Diploid | 1 | 1 |
| NL_0982 | swab_0982 | Buccal swab | Netherlands | Valthe | 52.8503 | 6.8827 | *P. lessonae* | *P.esculentus* | Diploid | 1 | 1 |
| NL_1604 | swab_1604 | Tissue | Netherlands | Visschersdijk | 52.2115 | 6.4758 | - | *P. lessonae* | Diploid | 2 | 0 |
| NL_1606 | swab_1606 | Tissue | Netherlands | Vragenderveen | 51.9798 | 6.6441 | - | *P.esculentus* | Diploid | 1 | 1 |
| NL_1607 | swab_1607 | Tissue | Netherlands | Vragenderveen | 51.9798 | 6.6441 | *P. lessonae* | *P. lessonae* | Diploid | 2 | 0 |
| NL_1580 | swab_1580 | Tissue | Netherlands | Wanninkhof | 52.1456 | 6.5114 | *P. lessonae* | *P. lessonae* | Diploid | 2 | 0 |
